# Distinct pattern of epigenetic DNA modification in leukocytes from patients with colorectal carcinoma and individuals with precancerous conditions, benign adenoma and inflammatory bowel disease – a link to oxidative stress

**DOI:** 10.1101/141903

**Authors:** Marta Starczak, Ewelina Zarakowska, Martyna Modrzejewska, Tomasz Dziaman, Anna Szpila, Kinga Linowiecka, Jolanta Guz, Justyna Szpotan, Maciej Gawronski, Anna Labejszo, Ariel Liebert, Zbigniew Banaszkiewicz, Maria Klopocka, Marek Foksinski, Daniel Gackowski, Ryszard Olinski

## Abstract

A characteristic feature of malignant cells, including colorectal cancer cells, is a profound decrease in level of 5-hydroxymethylcytosine, product of 5-methylcytosine oxidation by TET enzymes. This study included four groups of subjects: healthy controls, and patients with inflammatory bowel disease (IBD), benign polyps and colorectal cancer. Patients from all groups presented with significantly lower levels of 5-methylcytosine and 5-hydroxymethylcytosine than the controls. A similar tendency was also observed for 5-hydroxymethyluracil level. Patients with IBD showed the highest levels of 5-formylcytosine and 8-oxo-7,8-dihydro-2’-deoxyguanosine of all study subjects, and individuals with colorectal cancer presented with the lowest concentrations of vitamin C and A. Expressions of TET1 and TET2 turned out to be the highest in IBD group. To the best of our knowledge, this is the first study to show that healthy subjects, individuals with precancerous conditions and colorectal cancer patients present with distinct specific patterns of epigenetic modifications in leukocyte DNA.

## Introduction

It is generally accepted that mutations and aberrant methylation patterns are early events and important determinants of colon carcinogenesis. However, differences in colon cancer incidence seem to primarily reflect influence of environmental factors, among them oxidative stress which may be also linked to epigenetic changes [1-4].

Methylation of cytosine, usually involving CpG dinucleotides, is a key epigenetic modification, closely linked to gene repression, a process that exerts a profound effect on cellular identity and organismal fate [5]. Equally important is active DNA demethylation, a recently discovered process resulting in activation of previously silenced genes. Molecular background of active DNA demethylation is still not fully understood (reviewed in [6]). The most plausible mechanism involves ten-eleven translocation (TET) proteins that catalyze oxidization of 5-methylcytosine (5-mCyt) to 5-hydroxymethylcytosine (5-hmCyt), and then to 5-formylcytosine (5-fCyt) which is eventually converted to 5-carboxycytosine (5-caCyt) [6,7]. Some evidence from experimental studies suggests that TET may be also involved in synthesis of 5- hydroxymethyluracil (5-hmUra), a compound with epigenetic function [8].

A plethora of recent studies clearly demonstrated that 5-hmCyt is profoundly reduced in many types of human malignancies, including colorectal cancer (CRC) [9-11]. Moreover, in all previous studies, 5-hmCyt level in malignant tissue was always lower than in matched non-malignant control specimens [9-11]. However, it is still unclear whether this phenomenon is limited solely to cancer tissue, or is also present in surrogate materials from cancer patients, for example, leukocytes.

Furthermore, we still do not know what is the reason behind and a mechanism of 5-hmCyt decrease in cancer patients. Perhaps, this may be a consequence of reduced activity/expression of TET proteins [12]. However, it cannot also be excluded that active DNA demethylation taking place under altered conditions or in a different environment, for example in presence of chronic inflammation (that may induce oxidative stress) or in a person with poor nutritional status (that may influence vitamin C level, see also below), may modulate TET activity and thus, affect the level of epigenetic modifications.

Noticeably, our previous studies demonstrated that CRC patients present with significantly (ca. 30%) reduced levels of vitamin C [13,14]. Recent studies showed that ascorbate may enhance generation of 5-hmCyt in cultured cells [15-18]. Also vitamin A has been recently demonstrated to enhance 5-hmCyt synthesis and to modulate TET level [19]. Consequently, it cannot be excluded that the level of epigenetic DNA modifications and the expression of TETs in leukocytes are associated with blood concentrations of vitamin C and A.

Although many previous studies centered around the determination 5-hmCyt level, only few authors analyzed 5-fCyt, 5-caCyt and 5-hmUra contents in various tissues [8,20-25]. Due to very low level of these modifications in mammalian genome (approximately 3-4 orders of magnitude lower than 5-hmCyt content), their accurate determination is highly challenging. In this study, we used our recently developed rapid, highly-sensitive and highly-specific isotope-dilution automated online two-dimensional ultra-performance liquid chromatography with tandem mass spectrometry (2D-UPLC-MS/MS) [26,27] to analyze global methylation and the levels of all the abovementioned TET-mediated oxidation products of 5-mCyt and thymine: 5-hmCyt, 5-fCyt, 5-caCyt and 5-hmUra. Moreover, we determined leukocyte content of the best characterized marker of oxidatively modified DNA, 8-oxo-7,8-dihydro-2’-deoxyguanosine (8-oxodG), as well as the expression of TETs mRNA, and plasma concentrations of ascorbate and retinol.

Leukocytes are often used as an easily accessible surrogate marker providing information about environmentally-induced DNA modifications in other tissues [28,29]. In our present study, we examined leukocytes from CRC patients and individuals with two conditions being the most common etiological factors of sporadic colorectal malignancies, colon polyps (AD) and inflammatory bowel disease (IBD). Since CRC, AD and IBD are associated with oxidative stress, we also analyzed the level of the best characterized marker of oxidatively modified DNA, 8-oxodG.

This study fills a gap in existing knowledge about characteristics of TET-mediated DNA modifications and oxidatively modified DNA in a surrogate tissue (leukocytes) from patients with various conditions predisposing to CRC.

## Results

### Levels of epigenetic modifications and 8-oxodG in leukocyte DNA

The highest levels of 5-mdC and 5-hmdC were found in healthy controls, followed by patients with IBD, AD and CRC (Table. 1 and Fig. 1A and 1B), with the lowest levels of 5-mdC and 5-hmdC observed in AD and CRC group, respectively. All patients, irrespective of their primary disease, presented with significantly lower levels of 5-mdC and 5-hmdC than the controls. Similar to 5-mdC and 5-hmdC, also the level of 5-hmdU turned out to be significantly higher in healthy persons than in all groups of patients (Table. 1 and Fig. 1E). Among patients, the highest level of 5-hmdU was observed in AD group and the lowest in CRC group; also this intergroup difference was statistically significant. Persons with IBD presented with significantly higher 5-fdC level than patients with other diseases and healthy controls (Fig. 1C and Table. 1). Also the level of 8-oxodG in IBD group turned out to be significantly (1.6- to 2.9-fold) higher than in other study groups (Fig. 1F and Table. 1). Patients with CRC presented with the lowest level of 8-oxodG and the highest level of 5-cadC of all the study subjects. In turn, the lowest level of the latter modification was found in AD patients (Fig. 1D and Table. 1). 5-cadC levels in patients with AD and IBD were significantly lower than in the controls.

**Table 1.** Descriptive statistics for analyzed parameters in healthy controls and patients with inflammatory bowel disease (IBD), colon polyps (AD) and colorectal cancer (CRC)

|  | control | IBD | polyp | cancer |
| --- | --- | --- | --- | --- |
|  | geometric mean<br>(min;max) | geometric mean<br>(min;max) | geometric mean<br>(min;max) | geometric mean<br>(min;max) |
|  | median<br>(interquartile range) | median<br>(interquartile range) | median<br>(interquartile range) | median<br>(interquartile range) |
| 5-methyl-2'-<br>deoxycytidine<br>per 10<sup>3</sup> dN | 8.914<br>(8.155;9.711) | 8.567<br>(7.655;9.232) | 8.422<br>(7.833;9.403) | 8.545<br>(7.005;9.858) |
|  | 8.934<br>(8.722-9.080) | 8.762<br>(8.286-8.853) | 8.401<br>(8.216-8.572) | 8.566<br>(8.300-8.786) |
| 5-(hydroxymethyl)-2'-<br>deoxycytidine<br>per 10<sup>3</sup> dN | 0.066<br>(0.036;0.098) | 0.052<br>(0.005;0.100) | 0.053<br>(0.017;0.099) | 0.051<br>(0.015;0.085) |
|  | 0.068<br>(0.060-0.076) | 0.060<br>(0.041-0.077) | 0.052<br>(0.044-0.066) | 0.051<br>(0.043-0.063) |
| 5-formyl-2'-deoxycytidine<br>per 10<sup>6</sup> dN | 0.082<br>(0.013;0.758) | 0.190<br>(0.048;12.075) | 0.105<br>(0.020;5.028) | 0.115<br>(0.026;5.640) |
|  | 0.081<br>(0.051-0.129) | 0.122<br>(0.104-0.183) | 0.094<br>(0.068-0.137) | 0.087<br>(0.060-0.136) |
| 5-carboxy-2'-<br>deoxycytidine<br>per 10<sup>9</sup> dN | 10.031<br>(2.679;43.221) | 8.814<br>(2.882;46.340) | 7.606<br>(2.855;77.192) | 13.330<br>(2.998;55.269) |
|  | 9.779<br>(6.060-15.460) | 7.364<br>(5.661-12.809) | 7.086<br>(4.538-9.997) | 12.411<br>(8.084-23.029) |
| 5-(hydroxymethyl)-2'-<br>deoxyuridine per 10<sup>6</sup> dN | 0.546<br>(0.041;6.759) | 0.367<br>(0.114;3.599) | 0.425<br>(0.062;4.159) | 0.302<br>(0.055;4.444) |
|  | 0.643<br>(0.169-1.564) | 0.304<br>(0.208-0.563) | 0.353<br>(0.267-0.544) | 0.286<br>(0.171-0.482) |
| 8-oxo-7,8-dihydro-2'-<br>deoxyguanosine<br>per 10<sup>6</sup> dN | 1.089<br>(0.309;3.886) | 2.662<br>(0.688;20.659) | 1.065<br>(0.144;3.902) | 0.843<br>(0.131;4.321) |
|  | 1.103<br>(0.697-1.714) | 2.594<br>(1.948-3.258) | 1.641<br>(0.305-2.692) | 0.891<br>(0.545-1.466) |
| plasma ascorbate<br>concentration [μM] | 35.698<br>(1.057;102.178) | 24.303<br>(0.345;117.367) | 23.568<br>(0.335;127.489) | 15.562<br>(0.383;111.076) |
|  | 42.525<br>(31.405-55.790) | 31.157<br>(14.761-43.942) | 31.533<br>(9.071-62.059) | 23.213<br>(5.664-44.138) |
| plasma retinol<br>concentration [μM] | 1.703<br>(0.780;3.067) | 1.493<br>(0.062;3.835) | 1.704<br>(0.339;4.189) | 1.084<br>(0.217;4.620) |
|  | 1.739<br>(1.418-2.074) | 1.803<br>(1.091-2.150) | 1.749<br>(1.457-2.169) | 1.150<br>(0.793-1.513) |
| TET1 mRNA expression<br>ratio | 0.031<br>(0.010;0.165) | 0.414<br>(0.010;17.200) | 0.108<br>(0.010;9.203) | 0.070<br>(0.013;6.798) |
|  | 0.027<br>(0.013-0.081) | 2.076<br>(0.024-3.632) | 0.090<br>(0.028-0.218) | 0.054<br>(0.018-0.190) |
| TET2 mRNA expression<br>ratio | 7.629<br>(0.000;112.000) | 36.386<br>(6.813;179.700) | 16.042<br>(2.489;82.270) | 15.916<br>(0.000;199.700) |
|  | 19.610<br>(11.470-38.750) | 38.920<br>(18.900-67.220) | 19.315<br>(6.660-37.520) | 29.150<br>(13.120-71.250) |
| TET3 mRNA expression<br>ratio | 0.500<br>(0.031;5.385) | 0.437<br>(0.000;12.850) | 0.329<br>(0.002;4.204) | 0.094<br>(0.000;38.230) |
|  | 0.435<br>(0.258-1.051) | 0.634<br>(0.304-1.131) | 0.547<br>(0.090-1.619) | 0.509<br>(0.143-1.103) |

**Fig. 1.**
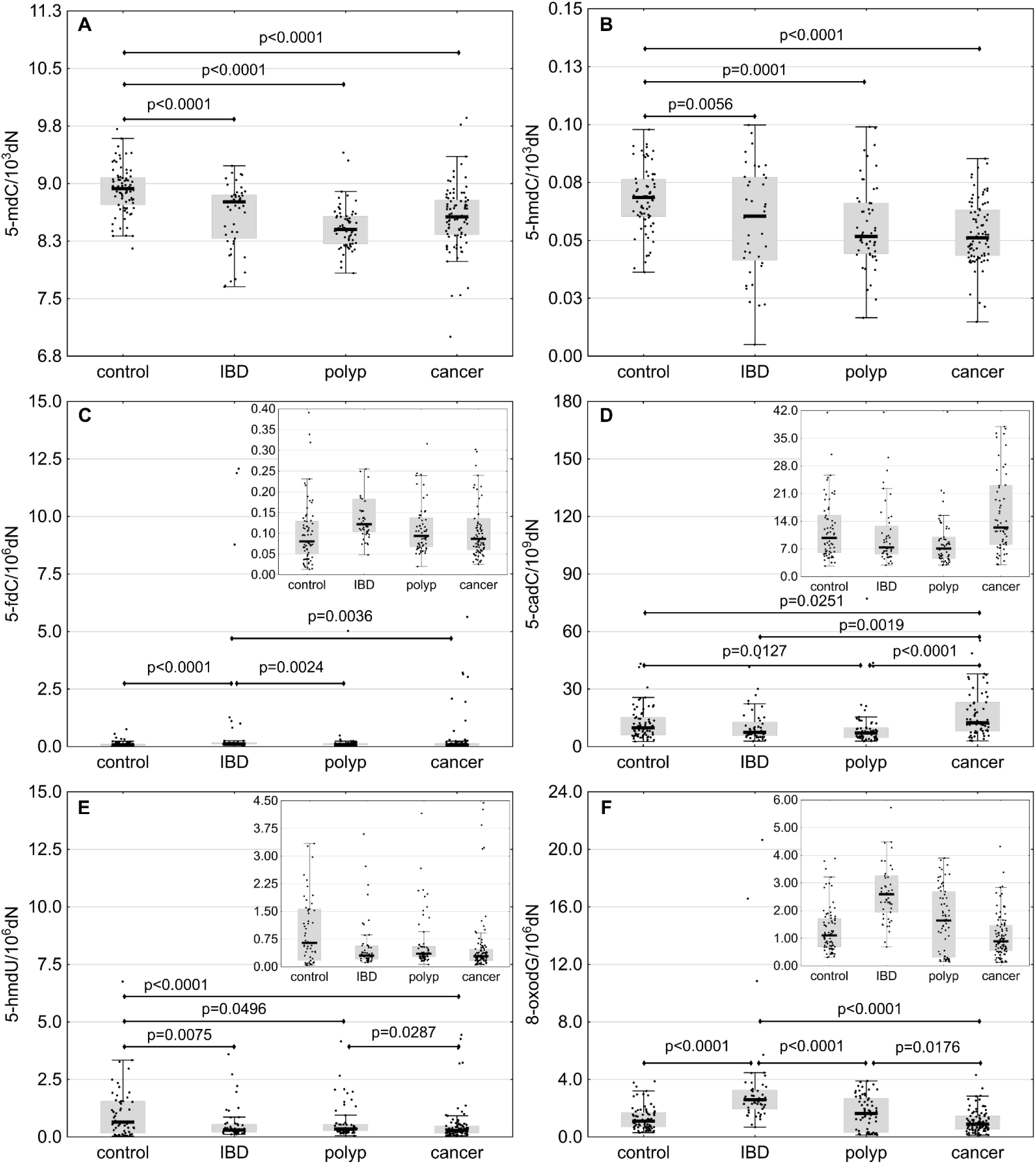
Levels of epigenetic modifications and 8-oxodG in leukocyte DNA from healthy controls and patients with IBD, AD (polyp) and CRC (cancer). The results presented as medians, interquartile ranges and non-outlier ranges. Raw (5-mdC) and normalized (other parameters) values were subjected to one-way analysis of variance (ANOVA) with LSD and Tukey post-hoc tests.

### Concentrations of vitamin C (ascorbic acid) and vitamin A (retinol) in blood plasma

Plasma concentration of ascorbic acid was the highest in healthy controls and the lowest in CRC patients (Fig. 2A and Table. 1). Individuals with CRC presented with significantly lower levels of vitamin C than other study subjects. Additionally, plasma concentration of ascorbate in persons with AD turned out to be significantly lower than in the controls. Plasma concentration of reti nolin patients with CRC was significantly lower than in other study subjects (Fig. 2B and Table. 1). In turn, the highest levels of retinol were found in IBD patients.

**Fig. 2.**
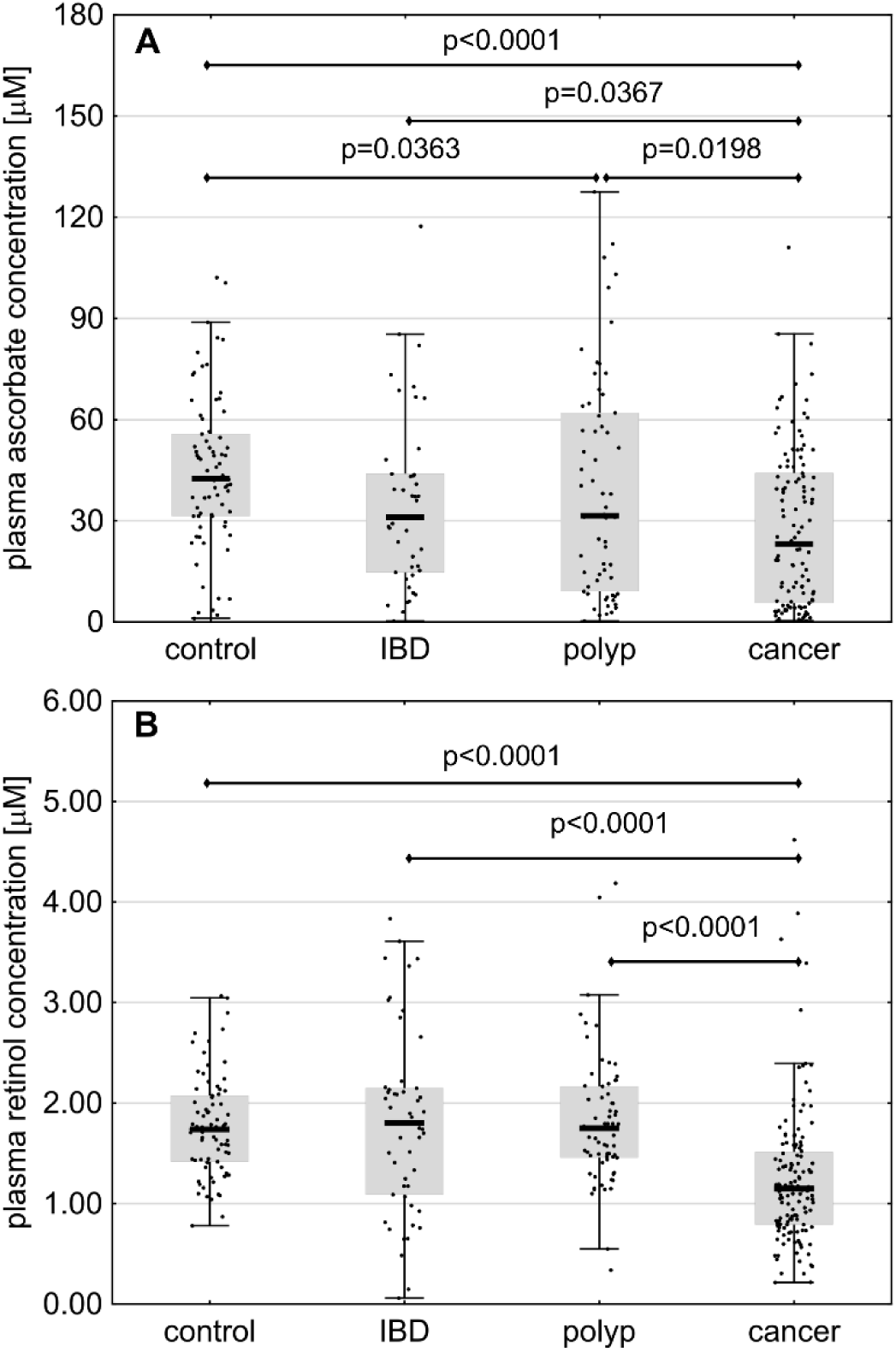
Plasma concentrations of vitamin C (ascorbic acid) and vitamin A (retinol) in healthy controls and patients with IBD, AD (polyp) and CRC (cancer). The results presented as medians, interquartile ranges and non-outlier ranges. Raw (reti nol) or normalized (ascorbic acid) values were subjected to one-way analysis of variance (ANOVA) with LSD and Tukey post-hoc tests.

### Expression of TET mRNA

Expression of TET1 in individuals with IBD turned out to be significantly higher than in other study subjects (Fig. 3A and Table. 1). Moreover, statistically significant differences were found in TET2 expressions in IBD patients and controls, as well as in IBD and AD groups (Fig. 3B and Table. 1). The study groups did not differ significantly in terms of TET3 expressions (Fig. 3C and Table. 1).

**Fig. 3.**
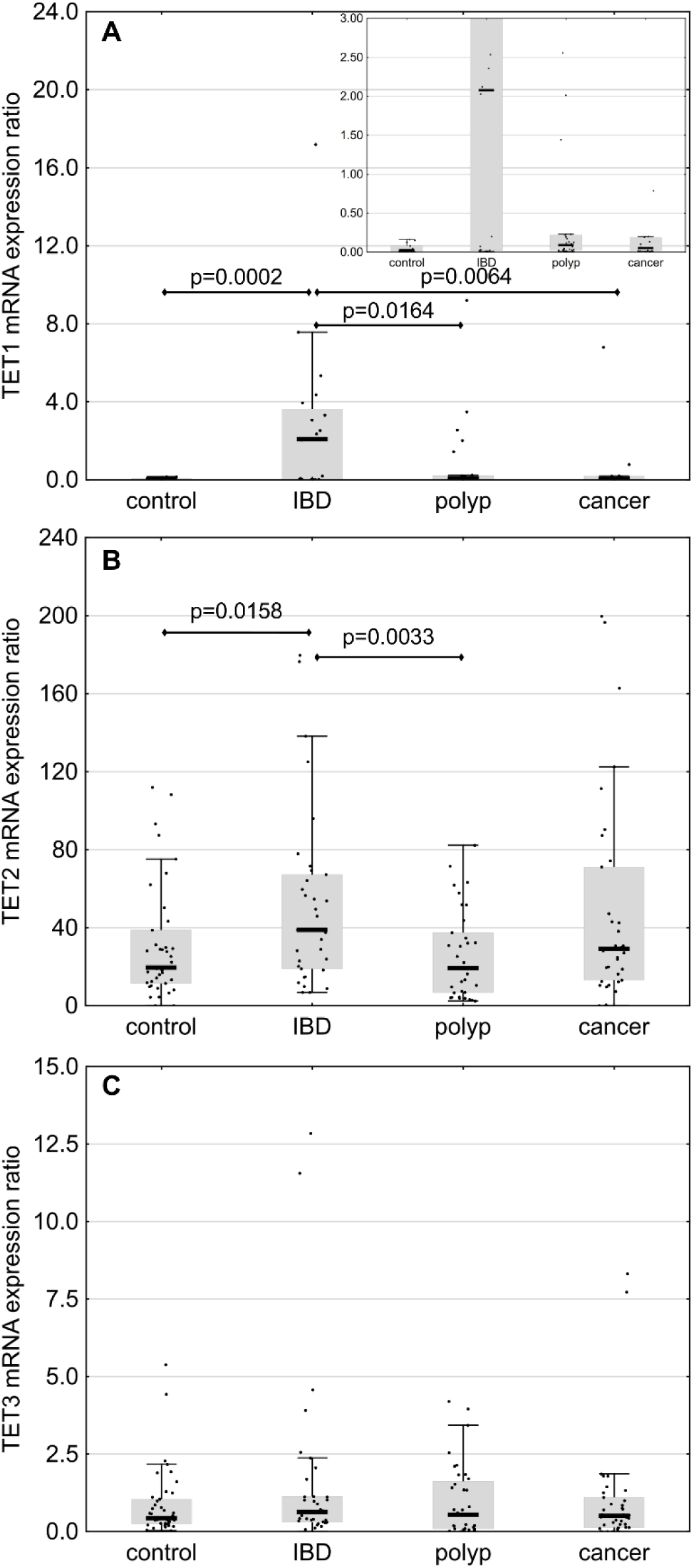
Expression of TET mRNA in healthy controls and patients with IBD, AD (polyp) and CRC (cancer). The results presented as medians, interquartile ranges and non-outlier ranges. Normalized values were subjected to one-way analysis of variance (ANOVA) with LSD and Tukey post-hoc tests.

### Correlations

Irrespective of the patients study group, the level of 8-oxodG correlated significantly with the levels of two epigenetic modifications, 5-fdC and 5-cadC. Furthermore, a significant correlation was found between 8-oxodG and 5-hmdU levels in IBD group. 8-oxodG level in AD group correlated positively with 5-hmdC and 5-hmdU levels; none of these correlations was observed in the controls. Healthy controls, as well as patients with AD and CRC, showed statistically significant substrate-product correlations between 5-fdC and its derivative, 5-cadC. A strong inverse correlation between 5-mdC and 8-oxodG was found in IBD patients, along with an inverse correlation between 5-mdC and 5-fdC. Moreover, 5-mdC level in persons with IBD correlated significantly with 5-hmdC content. In patients with AD, 5-hmdU showed significant positive correlations with all other epigenetic modifications, 5-hmdC, 5-fdC and 5-cadC. In turn, 5-hmdU level in CRC patients and controls correlated significantly solely with 5-fdC level. Finally, statistically significant correlations were found between 5-mdC and 5-cadC levels in healthy controls, as well as between 5-hmdC and 5-cadC levels in both healthy controls and CRC patients (Figure S1-S5).

In AD group, statistically significant positive correlations were found between plasma concentrations of ascorbic acid and 5-hmdC, 5-hmdU and 8-oxodG levels in leukocyte DNA. Individuals with plasma concentrations of vitamin C below the lower and above the upper quartile differed significantly in terms of their 5-hmdC levels (Fig. 4A). Plasma ascorbate concentration in the controls correlated positively with 5-hmdU level.

**Fig. 4.**
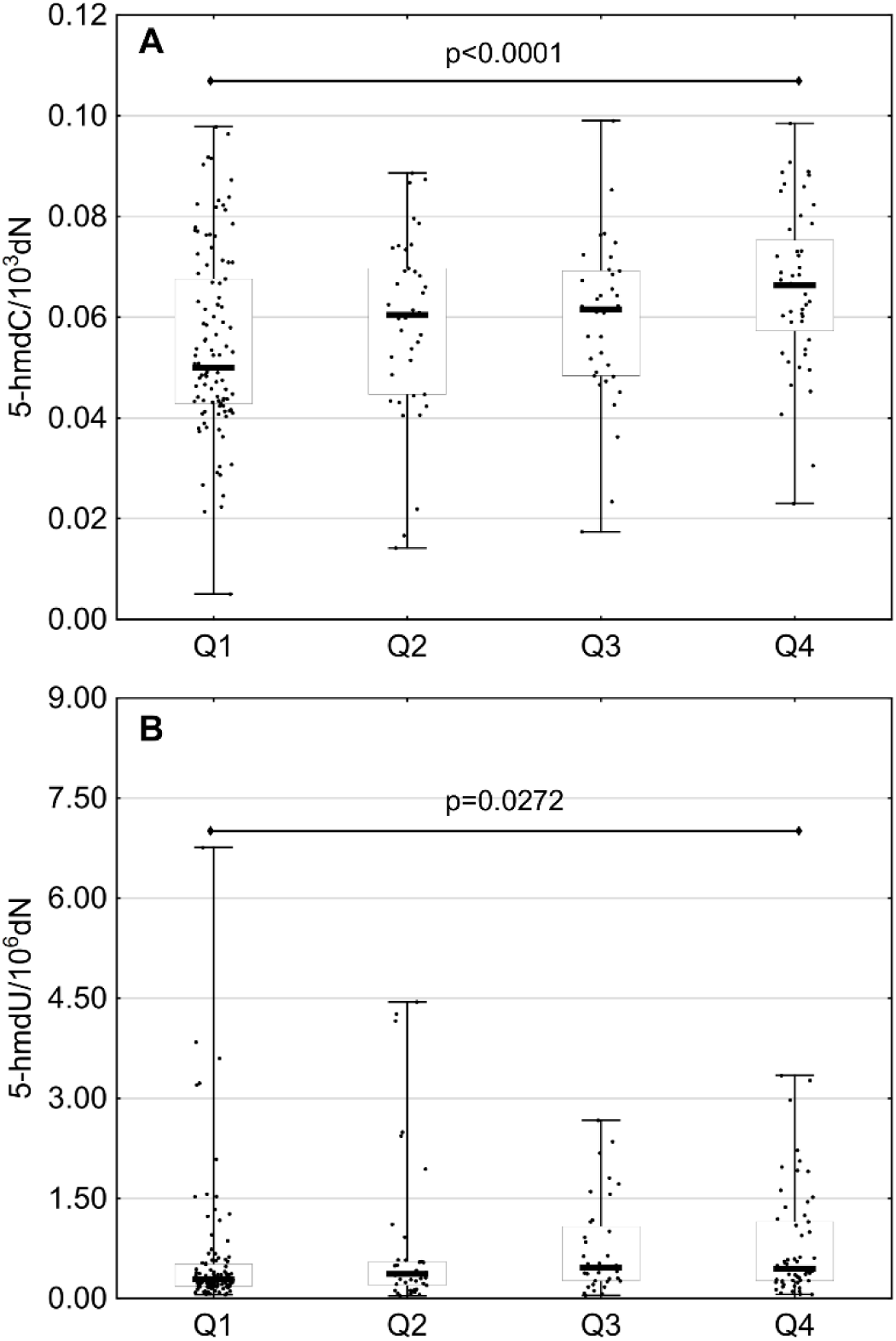
Selected, statistically significant associations between the levels of DNA modifications and plasma concentrations of vitamin C. Complete correlation matrix is presented in Supplementary Information.

A significant positive correlation was found between TET2 and TET1 expressions. Statistically significant inverse correlations were observed between the expression of TET2 and the levels 5-fdC and 5-hmdU in IBD group, along with the inverse correlations between TET3 expression, 5-hmdC and 5-hmdU levels. Finally, statistically significant inverse correlations were found between TET1 expression and 5-hmdU level in AD group, and between TET2 expression and 5-cadC level in patients with CRC (Figure S1-S5).

## Discussion

CRC usually develops as a consequence of neoplastic transformation of colon epithelial cells; usually, the first stage of colonic carcinogenesis is a benign polyp that may eventually progress to an invasive carcinoma. Thus, the group of AD patients analyzed in this study might include some proportion of individuals at early stages of colon cancer development. Also individuals with IBD are at increased risk of CRC [30]. The risk of CRC increases gradually with the time elapsed since the onset of IBD, which implies that carcinogenesis may be promoted by chronic inflammation. However, to the best of our knowledge, none of the previous studies analyzed an association between chronic inflammation and generation of 5-hmCyt derivatives in leukocytes. This is quite surprising owing that some metabolic changes associated with inflammation may influence formation of 5-hmCyt and its derivatives.

Although a molecular link between colon adenomas/IBD and carcinogenesis is yet to be established, it likely involves aberrant methylation and oxidative damage of DNA (reviewed [30,31]), and these processes were postulated to precede colonic dysplasia and CRC development [32]. Furthermore, it cannot be excluded that aberrant methylation of DNA is somehow related to oxidative stress. A growing body of evidence suggests that reduced content of 5-hmCyt may be characteristic not only for cancer tissue but also for precancerous lesions [11]. This in turn suggests that this process may continuously potentiate during tumor progression.

Our present study showed for the first time that 5-hmCyt content in leukocytes decreased gradually according to the following pattern: healthy controls > IBD patients > AD patients > CRC patients (Fig. 1B). This suggests that a decrease in global level of 5-hmCyt observed during the course of colon cancer development is not limited solely to the malignant tissue, but may also occur in surrogate tissues, such as leukocytes. This in turn implies that aberrant methylation of DNA may be a systemic process, rather than a local phenomenon. Aside from a decrease in 5-hmCyt content, leukocytes from all patient groups contained significantly less 5-hmUra than the cells from healthy controls (Fig. 1E).

As already mentioned, recent evidence suggests that TETs may catalyze synthesis of 5-hmUra from thymine. According to Pfaffender et al. [8], 5-hmUra level changes during the course of epigenetic reprogramming of the cell, following the same pattern as other products of TETs, i.e. 5-hmCyt, 5-caCyt and 5-fCyt. This implies that 5-hmUra may have an epigenetic function, similar to other products of active DNA demethylation. Moreover, 5-hmUra present in DNA was shown to recruit transcription factors and proteins involved in chromatin remodeling [8]. Our hereby presented results add to this evidence, suggesting that similar to 5-hmCyt, also 5-hmUra may be an epigenetic mark of carcinogenesis.

The level of 5-fCyt, a higher-order oxidative epigenetic mark, turned out to be significantly higher in IBD patients than in other groups analyzed in our study (Fig. 1C). Interestingly, individuals with IBD presented also with significantly higher level of 8-oxodG, an established marker of oxidative stress (Fig. 1F).

The association between inflammation and oxidative stress is well documented, and a number of previous studies demonstrated that inflammatory conditions and infections may be associated with an increase in 8-oxo-dG levels. Inflammatory response may result in recruitment of activated leukocytes. This may lead to a “respiratory burst”, i.e. an increase in oxygen uptake with resultant enhanced release of reactive oxygen species (ROS), such as superoxide and hydrogen peroxide, which eventually results in DNA damage (for review see [33]).

Since generation of 5-fCyt can be potentially mediated by ROS [34,35], an increase in the level of this modification observed in our study might be associated with oxidative stress, a characteristic feature of leukocytes from IBD patients. However, individuals with IBD did not differ from other groups in terms of their levels of 5-hmUra, a modification that is generally considered a product of free radical-mediated reaction with thymine [35]. Noticeably, all patients participating in this study, irrespective of their underlying condition, showed statistically significant positive correlations between the leukocyte levels of all analyzed oxidized epigenetic modifications (except 5-hmdC in IBD group) and 8-oxodG content, while no such associations were found in healthy controls (Figure S2-S5).

Endogenous synthesis of free radicals probably does not constitute a principal reason behind formation of epigenetic marks in cellular DNA [36]. Rather, environment characteristic for oxidative stress linked with the pathogenesis may influence factors responsible for the formation of abovementioned modifications. Indeed, recent evidence suggests that oxidative stress may contribute to post-translational modulation of TET2 [37]. In line with these findings, we showed that expression of TET1 and TET2 mRNA in leukocytes form IBD patients (characterized by the most severe oxidative stress) was significantly higher than in other study subjects (Fig. 3A and 3B). Furthermore, individuals with IBD presented with elevated levels of 5-fCyt. Since the structure of co-substrates for TETs (2-ketoglutarate, Fe^+2^) depends on redox state of the cell, the change in the activity of these enzymes may reflect the level of oxidative stress in IBD patients, i.e. the factor that might contribute to 5-fCyt formation. Moreover, it cannot be excluded that also superoxide (O ^−2^), an anion radical of dioxygen and the precursor of free radicals, plays an important role in TET-mediated active DNA demethylation [38,39].

The level of another higher-order oxidative modification of 5-mCyt, i.e. 5-caCyt, was the lowest in leukocytes from patients with precancerous conditions, AD and IBD (Fig. 1D). Recent evidence suggests that TET2 may yield 5-fCyt and 5-caCyt without the release and dilution of 5-hmCyt, and consecutive steps of iterative oxidation were postulated to be regulated by co-substrate levels [40]. Consequently, persistent increase in oxidative stress may alter TET activity, promoting/down-regulating generation of 5-fCyt and 5-caCyt during iterative oxidation of 5-mCyt (see also below). All this evidence suggests that synthesis of epigenetic DNA modifications is linked to oxidative stress; however, character of this relationship is complex and still not fully understood.

In our present study, patients from all groups presented with significantly lower levels of 5-mCyt than the controls (Fig. 1A), with the most evident decrease observed in individuals with AD and CRC. The distribution of 5-mCyt values across the study groups followed a similar pattern as for 5-hmCyt, i.e. healthy controls > IBD patients > AD and CRC patients. A dramatic decrease in both 5-mCyt and 5-hmCyt level may contribute to genomic instability, constituting a decisive step in CRC development. Interestingly, we found a significant inverse correlation between 5-mCyt and 8-oxodG levels in IBD patients (Fig. S3), which constitutes another argument for a potential link of aberrant DNA methylation to oxidative stress.

A few previous studies demonstrated that ascorbate may enhance generation of 5-hmCyt in cultured cells, probably acting as a cofactor of TETs during hydroxylation of 5-mCyt [15-18]. Recently, we have reported a spectacular increase in 5-hmUra content resulting from stimulation with vitamin C [16]. In turn, our present study demonstrated a positive correlation between plasma concentration of vitamin C and the levels of two epigenetic modifications, 5-hmCyt and 5-hmUra, in leukocyte DNA (Figure S1). Moreover, we found a significant difference in the levels of these modifications in patients whose plasma concentrations of vitamin C were below the lower and above the upper quartile (Fig. 4A and 4B). Probably, this is the first *in vivo* evidence of vitamin C involvement in generation of epigenetic DNA modifications. Our hereby presented findings suggest that ascorbate may play a role in cancer control, preventing aberrant methylation of DNA.

To the best of our knowledge, this is the first study to show that each of the analyzed groups, healthy controls, individuals with precancerous conditions and CRC patients, presented with a characteristic pattern of epigenetic modifications in leukocyte DNA. Therefore, an important question arises about the mechanism(s) involved in the development of epigenetic modification profiles being specific for a given setting? Perhaps, these were consequences of oxidative stress (differences in redox status) accompanying a given pathological condition, which contributed to alterations of cellular metabolism and interfered with iterative-enzymatic DNA modification.

While involvement of TETs in formation of all epigenetic modifications analyzed in this study raises no controversies, still little is known about the regulation of this process. Specifically, it is unclear why oxidation of 5-mCyt either stops at 5-hmCyt stage or proceeds to 5-fCyt and 5-caCyt stages. One potential explanation is different affinity of TETs to 5-mCyt, 5-hmCyt and 5-fCyt (for review see [41,42]). It is also possible that different proteins/factors recognize the modifications and determine their fate [43]. Interestingly, some recent experiments demonstrated that transcription factors, Myc and Max, and perhaps also a number of other regulatory proteins, may specifically recognize 5-caCyt, but have lesser affinity to 5-fCyt, and show only a trace of affinity to 5-mCyt and 5-hmCyt [44]. Moreover, a recent study conducted by Xiong et al. [45] showed that Sall4, an oncogenic protein being overexpressed in colon cancer [46], may further enhance TET2-catalyzed oxidation of 5-hmCyt.

To summarize, this study showed that environmental factors associated with some pathophysiological conditions linked to CRC development may alter the pattern of epigenetic modification, and thus, are involved in colorectal carcinogenesis. If supported by transcriptional information, the analysis of global epigenetic modification patterns in DNA from easily accessible cells, such as peripheral blood leukocytes, may provide a better insight into the mechanisms of carcinogenesis.

## Materials and methods

### Study group

The study included four groups of subjects: 1) healthy controls (n=79, median age 55 years, 63% of women), 2) patients with IBD (n=51, median age 35 years, 53% of women), 3) persons with colorectal polyps (AD), i.e. histologically confirmed adenoma tubulare (90%) or adenoma tubulovillosum (10%) (n=67, median age 65 years, 46% of women), and 4) individuals with colorectal cancer (CRC), i.e. histologically confirmed stage A (8%), stage B (45%), stage C (29%) or stage D (9%) adenocarcinoma, or malignant polyps (9%) (n=136, median age 65 years, 46% of women). None of the study subjects were related with one another, and all of them were Caucasians. All participants of the study were recruited in a hospital setting (Collegium Medicum, Nicolaus Copernicus University, Bydgoszcz, Poland) and subjected to colonoscopy. The study groups were matched for dietary habits, body weight and smoking status. The protocol of the study was approved by the Local Bioethics Committee, Collegium Medicum in Bydgoszcz, Nicolaus Copernicus University in Torun (Poland), and written informed consent was sought from all the participants.

### Isolation of DNA and determination of epigenetic modifications and 8-oxodG in DNA isolates

Leukocytes were isolated from heparinized blood samples with Histopaque 1119 (Sigma, United Kingdom) solution, according to the manufacturer’s instruction, and stored at -80°C until the analysis. Isolation of leukocyte DNA and its hydrolysis were carried out as previously described [27], but the cell pellet was immediately dispersed in ice-cold lysing buffer B, without homogenization and washing steps. Procedures used for the determination of 5-methyl-2’-deoxycytidine (5-mdC), 5-(hydroxymethyl)-2’-deoxycytidine (5-hmdC), 5-formyl-2’-deoxycytidine (5-fdC), 5-carboxy-2’deoxycytidine (5-cadC), 5-(hydroxymethyl)-2’-deoxyuridine (5-hmdU) and 8-oxodG by 2D-UPLC-MS/MS have been described elsewhere [27].

### Determination of vitamin C (ascorbic acid) in blood plasma by UPLC-UV

#### Sample preparation

To stabilize vitamin C (ascorbic acid) and to precipitate proteins, 200-μL aliquots of freshly prepared or partially thawed plasma were mixed with 200 μL of precooled 10% (w/v) meta-phosphoric acid (MPA, Sigma-Aldrich, Munich, Germany) containing uracil (50 μM) (Sigma-Aldrich, Munich, Germany) as an internal standard. The samples were kept on ice for 40 min and then diluted with 200 μL of MilliQ-grade deionized water (Merck Millipore, Darmstadt, Germany), vortexed and centrifuged at 25155×g for 15 min at 4°C. The supernatants (200 μL) were purified by ultrafiltration using AcroPrep Advance 96-Well Filter Plates 10K (Pall, Port Washington,New York, USA), and injected into Waters Acquity (Milford, Massachustts, USA) ultra-performance liquid chromatographic (UPLC) system. The method was validated with the reference material from Chromsystems (Gräfelfing, Germany).

#### Chromatography

The UPLC system consisted of binary solvent manager, sample manager, column manager and photo-diode array detector, all from Waters. The samples were separated on Waters Acquity UPLC HSS T3 column (150 mm×2.1 mm, 1.8 μm) with Van Guard HSS T3 1.8-μm pre-column at a flow rate 0.25 mL/min and 2-μL injection volume. Ammonium formate (10 mM, pH 3.1, Fluka, Munich, Germany) and acetonitrile (Sigma-Aldrich, Munich, Germany) were used as Solvent A and B, respectively. The following program was used for ascorbate elution: 0– 0.1 min 99% A, 1% B, 0.1–2.2 min 97% A, 2.2–4.0 min – linear gradient to 90% A, 4.0–4.5 min – 90% A, 4.5–6.0 min – 99% A. Column thermostat was set at 10°C. The effluent was monitored with a photo-diode array detector at 245 nm, and analyzed with Empower software.

### Determination of vitamin A (retinol) in blood plasma by HPLC-FD

#### Sample preparation

To precipitate proteins, 200-μL aliquots of freshly thawed plasma were mixed with 200 μL of anhydrous ethanol (POCH, Gliwice, Poland) containing an internal standard (0.5 ml IS/10 ml EtOH) (Vitamins A and E in Serum/Plasma – HPLC, Chromsystems, Gräfelfing, Germany), vortexed and left for 10-15 min. Then, 400 μL MilliQ-grade deionized water (Merck Millipore, Darmstadt, Germany) and 800 μL of hexane (Aldrich, Germany) were added to extract the vitamin. The samples were shaken vigorously for 2 h and centrifuged at 25155×g for 10 min. Then, 400 μL of the upper layer (hexane) were collected, dried in Speed-Vac system (8 min), and dissolved in 100 μL of mobile phase (acetonitrile and methanol (Sigma-Aldrich, Steinheim, Germany), 80:20 (v/v), HPLC-grade). The samples were shaken vigorously overnight, centrifuged at 25155×g for 10 min, and supernatants were injected into HPLC system. The method was validated with the reference material from Chromsystems (Gräfelfing, Germany).

#### Chromatography

The HPLC system consisted of 1525μ binary HPLC pump and 2707 autosampler, both from Waters. Vitamin A (retinol) was quantified with Jasco FP-920 fluorimetric detector (Jasco Co., Tokyo, Japan). The samples were separated in an isocratic system with a 5-μm Atlantis dC18 column equipped with guard cartridge (5 μm, 150 mm x 3.0, Waters). The mobile phase, consisting of acetonitrile methanol, 80:20 (v/v), was added at a flow rate of 1.5 ml/min, with 5-μL injection volume. The effluent was monitored by means of fluorimetric detection (*λ*_ex._=340 nm, *λ*_em._=472 nm for retinol, and *λ*_ex._=290 nm, *λ*_em._=330 nm for internal standard), and analyzed with Empower Software.

### Gene expression analysis

Isolated leukocytes were stored at −80°C until the analysis. RNA was isolated with MagNA Pure 2.0 (Roche) following the standard procedures. Concentration and purity of RNA aliquots were verified spectrophotometrically with NanoDrop 2000 (Thermo Scientific). A_260_/A_280_ ratio was used as an indicator of protein contamination, and A_260_/A_230_ ratio as a measure of contamination with polysaccharides, phenol and/or chaotropic salts. Quality and integrity of total RNA were assessed by visualization of 28S/18S/5.8*S rRNA* band pattern in a 1.2% agarose gel. Non-denaturing electrophoresis was carried out at 95 V for 20 min in TBE buffer (Tris – Boric Acid – EDTA). The gel was stained with ethidium bromide or SimplySafe and visualized using GBox EF Gel Documentation System (SynGene). Purified RNA was stored at −80°C. The samples with RNA concentrations greater than 50 ng/μl were qualified for further analysis. 0.5 microgram of total RNA from each sample (in 20-μl volume) was used for cDNA synthesis by reverse transcription with High-Capacity cDNA Reverse Transcription Kit (Applied Biosystems), according to the manufacturer’s instruction. The reaction was carried out with Mastercycler Nexus Gradient thermocycler (Eppendorf). To exclude contamination with genomic DNA, reverse transcriptase reaction included also a negative control. cDNA was either used for qPCR setup immediately after obtaining, or stored at −20°C.

The RT-qPCR complies with the Minimum Information for Publication of Quantitative Real-time PCR Experiments (MIQE) guidelines. Three gene transcripts, *TET1, TET2* and *TET3*, were analyzed by relative quantitative RT-PCR (RT-qPCR) with relevant primers and short hydrolysis probes substituted with Locked Nucleic Acids from the Universal Probe Library (UPL, Roche) (see: Tab. 2). The probes were labeled with fluorescein (FAM) at the 5’-end and with a dark quencher dye at the 3’-end. Expressions of target genes were normalized for two selected reference genes, *HMBS* (GeneID: 3145) and *TBP* (GeneID: 6908), using UPL Ready Assay #100092149 and #100092158, respectively. Real-time PCR mixes (in 20-μl volumes) were prepared from cDNA following the standard procedures for LightCycler480 Probes Master (Roche), provided with the reagent set. The reactions were carried out on 96-well plates. Aside from the proper samples, each plate included also no-template control and no-RT control. Quantitative real-time PCR was carried out with LightCycler 480 II, using the following cycling parameters: 10 s at 95°C, followed by 45 repeats, 10 s each, at 95°C, 30 s at 58°C, and finally, 1 s at 72°C with acquisition mode (parameters of wavelength excitation and detection equal 465 nm and 510 nm, respectively). The reaction for each gene was standardized against a standard curve, to estimate amplification efficiency. Standardization procedure included preparation of 10-fold serial dilutions with controlled relative amount of targeted template. The efficiency of amplification was assessed based on a slope of the standard curve. Standard dilutions were amplified in separate wells, but within the same run. Then, the samples were subjected to qPCR with measurement of *C*_t_, and amplification efficiencies were automatically calculated and displayed on the analysis window of LightCycler 480 software, version 1.5.1.62 (Roche). The same software was also used for sample setup, real-time PCR analysis and calculation of relative *C*_t_ values referred to as “Ratios”.

**Tab. 2.**
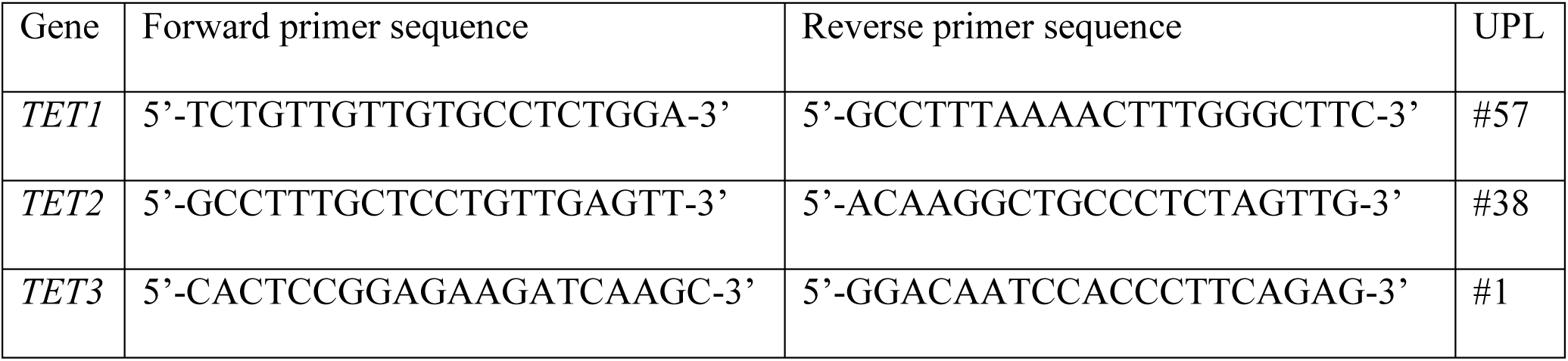
Primers and short hydrolysis probes used for TETs mRNA expression analysis.

### Statistical analysis

The results are presented as medians, interquartile ranges and non-outlier ranges. Normal distribution of the study variables was verified with Kolmogorov-Smirnov test with Lilliefors correction, and based on visual inspection of plotted histograms. Variables with non-normal distributions (5-hmdC, 5-fdC, 5-cadC, 5-hmdU, 8-oxodG, ascorbic acid concentration and TET mRNA expression) were subjected to Box-Cox transformation prior to statistical analyses with parametric tests. Normalized data were subjected to one-way analysis of variance (ANOVA) followed by LSD and Tukey post-hoc tests. Associations between pairs of variables were assessed based on Pearson correlation coefficients for raw or normalized data, where applicable. To assess the influence of ascorbate and retinol concentrations on the level of DNA modifications, the study subjects were divided into four subgroups (Q1-Q4) on the basis of the cut-off values corresponding to lower quartile, median and upper quartile for the control group. All statistical transformations and analyses were carried out with STATISTICA 13.1 PL [Dell Inc. (2016). Dell Statistica (data analysis software system), version 13. software.dell.com.]. The results were considered statistically significant at *P* values lower than 0.05.

## Acknowledgements

Funding: This work was supported by the Polish National Science Center [grant no. 2013/09/B/NZ5/00767].

---

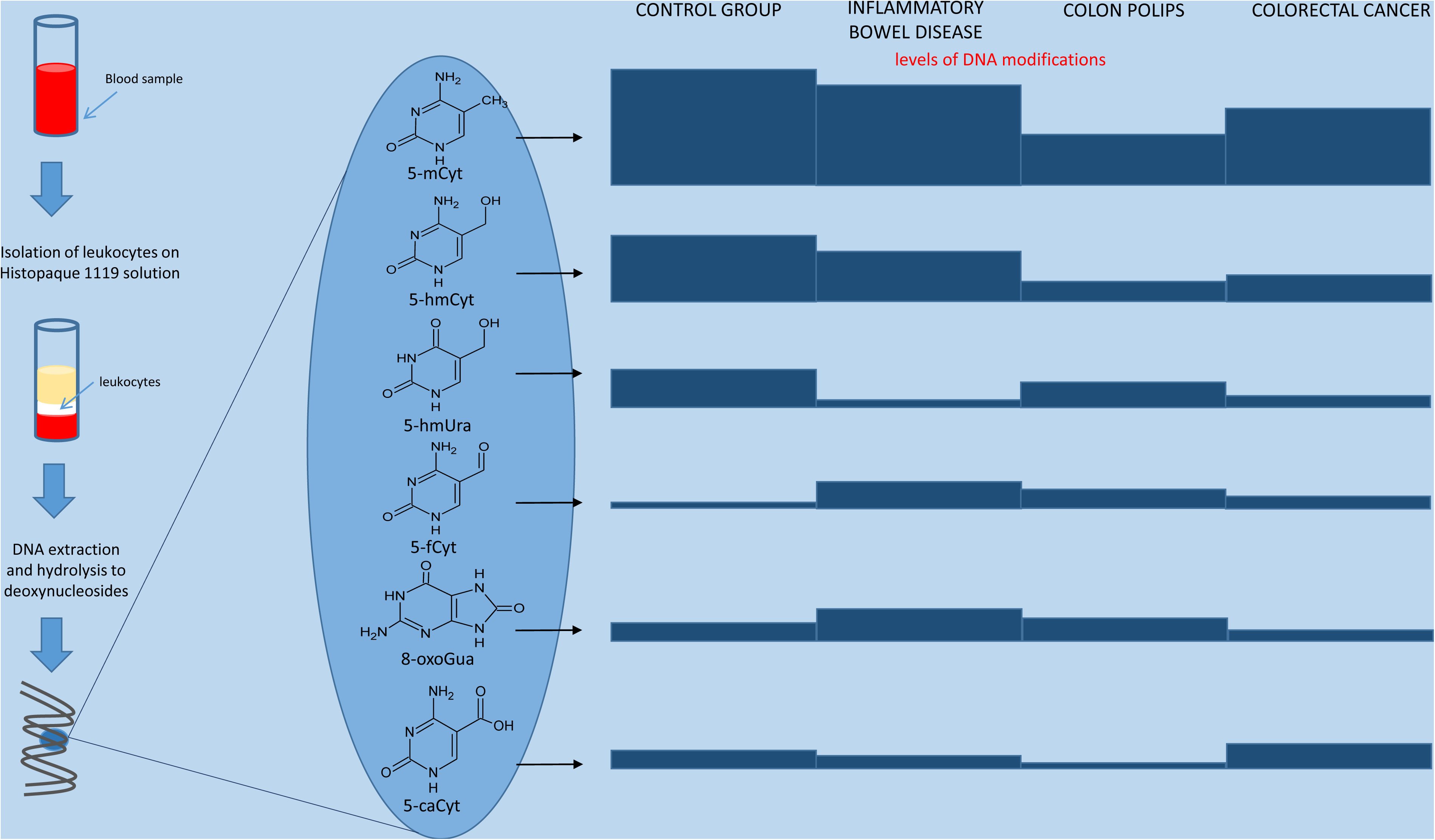

